# Repeated intermittent clenbuterol administration induces exposure history-dependent proteomic remodeling in mouse skeletal muscle

**DOI:** 10.64898/2026.07.24.740506

**Authors:** Keisuke Hitachi, Hisateru Yamaguchi, Yuri Kiyofuji, Kunihiro Tsuchida

**Affiliations:** Division for Therapies against Intractable Diseases, Center for Medical Science, Fujita Health University, Toyoake, Japan; Department of Management Nutrition, Aichi Gakusen University, Okazaki, Japan; Department of Biomedical Molecular Sciences, Fujita Health University School of Medicine, Toyoake, Japan

**Keywords:** clenbuterol, skeletal muscle, β2-adrenergic stimulation, proteomics, muscle hypertrophy, repeated exposure

## Abstract

Clenbuterol (CB), a β2-adrenergic receptor agonist, is known to increase skeletal muscle mass. However, because its effects are influenced by treatment duration, dose, and β-adrenergic receptor responsiveness, it remains unclear how extended intermittent CB exposure alters the skeletal muscle proteome. In this study, we established a repeated intermittent CB administration model in mice and compared the effects of single and repeated CB exposure on plantaris muscle using label-free quantitative proteomics. Repeated CB exposure produced a more evident muscle-weight response than single exposure under the same long-term experimental timeline. Proteomic profiling revealed that repeated CB exposure was not a simple reproduction or amplification of the single CB response, but was associated with distinct changes in sarcomere-associated, tissue-remodeling-related, membrane-trafficking-related, and metabolic proteins. Integrated response-class analysis further identified multiple patterns of protein regulation, including shared CB-responsive proteins and proteins preferentially identified as changed under repeated CB conditions. Immunoblotting validated representative proteins from these response classes: Klhl40 and Napa as shared CB-responsive proteins, Galectin-3 as a strongly CB-responsive remodeling-related protein, and Rab18 as a protein change more clearly detected under repeated CB conditions. Together, these findings indicate that repeated intermittent CB exposure induces exposure history-dependent proteomic remodeling in skeletal muscle. This study provides a resource for understanding the molecular consequences of repeated CB exposure and for future mechanistic studies of β2-adrenergic stimulation-induced skeletal muscle adaptation.

## Introduction

Skeletal muscle is a highly plastic tissue that adapts its mass and functional properties in response to physical activity, nutritional status, neural activity, endocrine stimuli, and pharmacological interventions. Among pharmacological stimuli, β2-adrenergic receptor stimulation has been widely studied because of its effects on skeletal muscle mass, metabolism, and body composition [1,2]. CB is a β2-adrenergic receptor agonist that has been extensively used as an experimental anabolic stimulus in skeletal muscle research. Animal studies have shown that CB administration increases skeletal muscle mass and can suppress muscle loss in models of muscle atrophy [3,4]. In addition, the hypertrophic and anti-atrophic effects of CB are preserved in β1-adrenergic receptor-deficient mice but abolished in β2-adrenergic receptor-deficient mice, demonstrating that the skeletal muscle effects of CB require β2-adrenergic receptors [4]. Consistent with the broader role of β2-adrenergic receptor signaling in skeletal muscle, other β2-adrenergic agonists, including formoterol and salmeterol, have also been reported to induce skeletal muscle hypertrophy in animal models [5]. More recently, chronic CB treatment has been shown to improve glucose metabolism in a skeletal muscle β2-adrenergic receptor- and Gs-dependent manner and to induce transcriptomic and metabolomic reprogramming in skeletal muscle cells [6]. Furthermore, β-arrestin 1 has been shown to be required for CB-induced skeletal muscle hypertrophy and changes in contractile properties in vivo as a downstream mechanism of β2-adrenergic receptor signaling [7].

The effects of CB are highly dependent on treatment conditions. Low-dose CB can induce skeletal muscle hypertrophy, whereas higher doses are associated with myotoxicity [8]. Chronic CB treatment has been reported to reduce β-adrenoceptor density in skeletal muscle [9]. In addition, CB decreases β2-adrenoceptor mRNA expression in fast-twitch fiber-enriched muscles, including the plantaris muscle, without altering its expression in the soleus muscle [10]. These changes in receptor responsiveness suggest that skeletal muscle responses to β2-adrenergic stimulation may depend not only on dose and treatment duration, but also on exposure history, including repeated exposure and recovery periods. Therefore, CB-induced skeletal muscle adaptation should be considered as a complex phenomenon shaped by dose, treatment duration, and prior exposure history, rather than simply as an acute or uniform hypertrophic response.

CB is also important from sports-science and social perspectives. The World Anti-Doping Agency lists CB as a prohibited anabolic agent and prohibits it at all times, both in- and out-of-competition (World Anti-Doping Code International Standard Prohibited List 2026). In addition, CB has been used illegally as a growth-promoting agent in livestock, and unintentional positive doping cases caused by contaminated meat have been reported [11,12]. Distinguishing intentional CB misuse from exposure through contaminated meat has also been examined as an important issue in sports doping [13]. In cases of CB misuse for fat loss or muscle gain, adverse events such as tachycardia, hypokalemia, electrocardiographic abnormalities, and rhabdomyolysis have been reported [14,15]. Thus, although CB has clear pharmacological effects on skeletal muscle, understanding the molecular changes caused by repeated exposure or misuse is important from both basic biological and anti-doping perspectives.

Skeletal muscle adaptation is established through molecular changes across multiple levels, including not only the accumulation and degradation of myofibrillar proteins, but also changes in fiber type, metabolic properties, extracellular matrix organization, and inflammatory and repair responses. Therefore, to understand the effects of β2-adrenergic receptor stimulation on skeletal muscle, it is important to capture global changes in protein expression profiles rather than focusing only on a single signaling pathway or representative markers of muscle hypertrophy. A previous study investigating CB-induced changes in the skeletal muscle proteome used rat plantaris muscle and showed, by histochemical analysis and 2D gel-based proteomic analysis, that low-dose, non-myotoxic CB treatment was associated with increased plantaris muscle weight, protein accretion, and preferential hypertrophy of fast oxidative glycolytic fibers [16]. In addition, a recent mass spectrometry (MS)-based proteomic analysis of human skeletal muscle reported that prolonged β2-adrenergic stimulation with terbutaline induces proteome remodeling that shares multiple features with resistance training, including changes in translational capacity-related proteins and increased abundance of kelch-like family proteins such as KLHL40 and KLHL41 [17]. However, few studies have directly compared single and repeated CB exposure within the same experimental system. It also remains unclear which protein responses are shared by both conditions and which become more evident after repeated exposure.

In this study, we established a repeated intermittent CB administration model in mice by combining 2-week treatment periods with 4-week washout periods. We then performed label-free quantitative proteomic analysis of plantaris muscles from control mice, mice exposed to a single period of CB treatment, and mice exposed to repeated CB treatment. Furthermore, by integrating three pairwise group comparisons, we performed response-class analysis to classify protein responses shared by single and repeated treatment and those associated with repeated exposure. Finally, representative differentially changed proteins identified by proteomic analysis were validated by immunoblotting. Our findings indicate that repeated intermittent CB exposure is not a simple reproduction of a single-exposure response, but is accompanied by exposure history-dependent proteomic responses in skeletal muscle.

## Materials and Methods

### Animals, experimental design, and muscle weight measurement

All mice were maintained in a temperature-controlled animal room at 25°C under a 12-h light/12-h dark cycle, with ad libitum access to chow and drinking water. Male C57BL/6NJ mice were obtained from the Jackson Laboratory Japan, Inc. and used from 9 weeks of age.

CB was supplied through the drinking water at 20 ppm (20 µg/mL) during each 2-week administration period. This concentration and treatment duration were based on a previous study [18]. Mice in the corresponding control periods received drinking water without CB. In the recovery experiment, mice were treated with or without CB in the drinking water for 2 weeks. Muscles were collected either immediately after the 2-week treatment period or after an additional 4-week period without CB. Control and CB-treated groups were included at both collection time points. For the repeated intermittent protocol, animals were divided into control, single CB, and repeated CB groups. The schedule comprised three 2-week treatment windows separated by 4-week intervals (Fig. 1A). Control mice received unsupplemented drinking water during all three windows. Single CB mice received unsupplemented drinking water during the first two windows and CB-containing water only during the final window. Repeated CB mice received CB-containing water during each of the three treatment windows. Animals were euthanized at the end of the final 2-week treatment period. Body weight was recorded immediately before tissue collection. Gastrocnemius (Gast), plantaris, and soleus muscles were excised, weighed immediately, snap-frozen in liquid nitrogen, and kept at −80°C until use.

**Figure 1.**
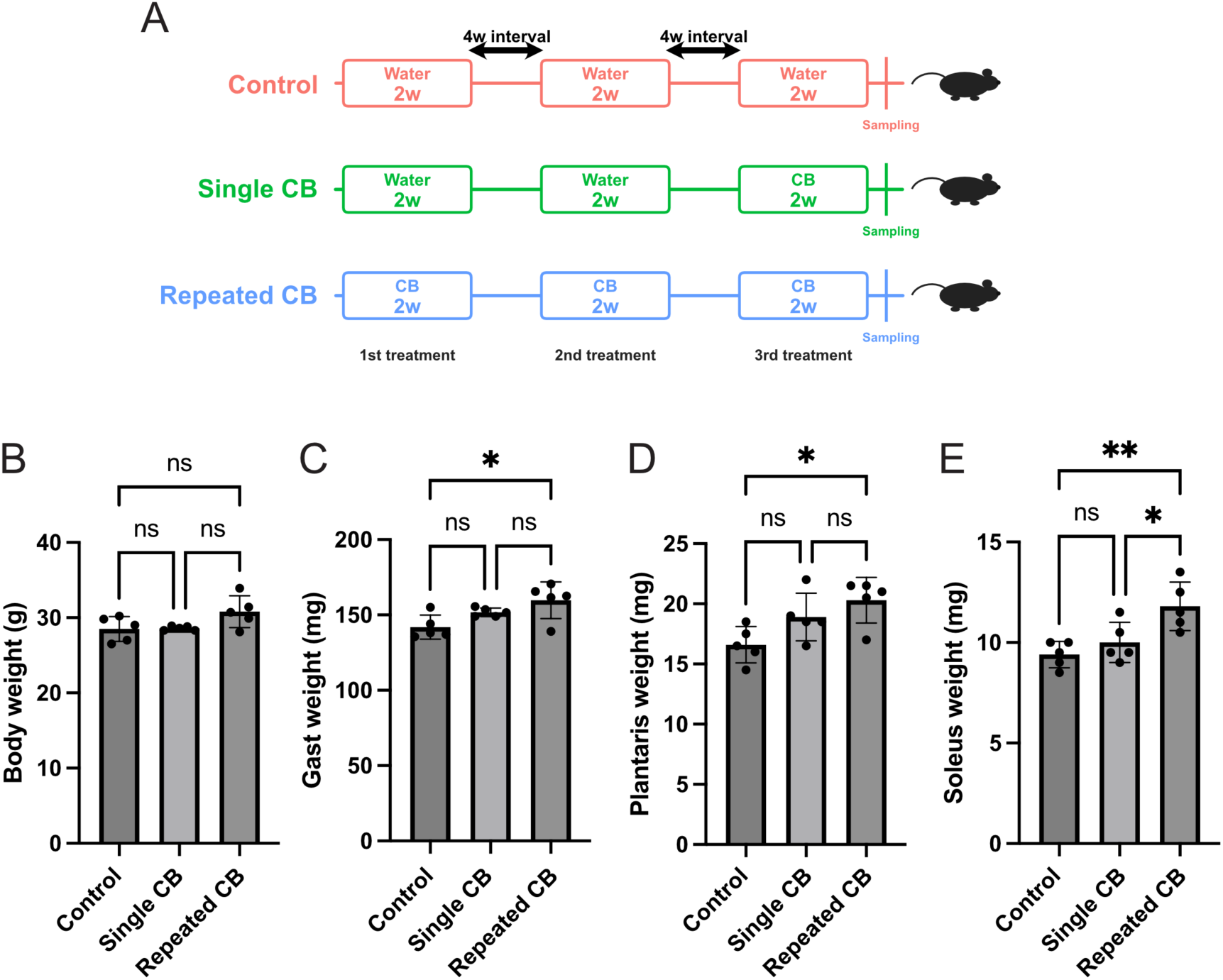
Repeated intermittent CB exposure produced a distinct muscle-weight response. (A) Experimental design of control, single CB, and repeated CB groups. The experimental timeline consisted of three 2-week treatment periods separated by 4-week intervals. Control mice received drinking water without CB during all three treatment periods. Single CB mice received drinking water without CB during the first and second treatment periods and CB-containing drinking water during the final treatment period. Repeated CB mice received CB-containing drinking water during all three treatment periods. Muscles were collected at the end of the final treatment period. (B–E) Body weight (B), Gast weight (C), plantaris weight (D), and soleus weight (E) in control, single CB, and repeated CB mice. Data are presented as mean ± s.d.; n = 5 mice per group. Data were analyzed by ordinary one-way ANOVA followed by Tukey’s multiple-comparison test. ns, not significant; *P < 0.05; **P < 0.01.

### Proteomic analysis

Plantaris muscles from the control, single CB, and repeated CB groups were processed for label-free proteomics. Frozen muscles were disrupted with a Shakeman 2 homogenizer (Bio Medical Science Inc., Japan) in RIPA buffer consisting of 50 mM Tris-HCl (pH 7.2), 150 mM NaCl, 0.1% SDS, 1% Triton X-100, and 0.5% sodium deoxycholate, supplemented with 4 µg/mL leupeptin, 1 mM PMSF, and 1 µg/mL aprotinin, based on previously used procedures [19,20]. Protein concentration was measured with the BCA Protein Assay Kit (Takara). Ten micrograms of total protein from each sample was subjected to SP3-based sample preparation [21]. Samples were reduced with dithiothreitol (DTT), alkylated with iodoacetamide (IAA), and digested with Trypsin/Lys-C Mix (Promega) at a 1:50 enzyme-to-substrate ratio. The resulting peptides were cleaned up with GL-Tip SDB tips (GL Sciences) before LC-MS/MS.

Peptide separation and MS acquisition were performed using an EASY-nLC 1000 nano-liquid chromatography system coupled to an Orbitrap Fusion ETD mass spectrometer (Thermo Fisher Scientific). Peptide samples were first trapped on an Acclaim PepMap 100 C18 column (3 µm, 75 µm ID × 20 mm; Thermo Fisher Scientific) and then separated on an EASY-Spray C18 analytical column (3 µm, 75 µm ID × 150 mm; Thermo Fisher Scientific). A 0–35% acetonitrile gradient in 0.1% formic acid was run at 300 nL/min. The spray voltage was 2 kV, and the ion transfer tube was maintained at 275°C. Xcalibur software (version 4.7.69.37; Thermo Fisher Scientific) was used for data-dependent acquisition. MS1 scans were recorded over an m/z range of 375–1500 at a resolution of 120,000. The most abundant precursor ions were subjected to collision-induced dissociation in the linear ion trap with a normalized collision energy of 35%, and dynamic exclusion was set to 60 s.

MS raw files were analyzed in Proteome Discoverer version 2.4.1.15 (Thermo Fisher Scientific) using SequestHT and Mascot (version 2.6.2). Spectra were searched against the Mus musculus UniProtKB database (TaxID = 10090; version 202104), to which an additional Myh2 entry (UniProt G3UW82) was added. Cysteine carbamidomethylation was used as a fixed modification, whereas methionine oxidation and protein N-terminal acetylation were used as variable modifications. The precursor and fragment ion tolerances were 10 ppm and 0.6 Da, respectively. Trypsin digestion was searched with a maximum of two missed cleavages. Peptide-spectrum matches were filtered by target-decoy searching at a 1% false discovery rate. Label-free abundance values were obtained in Proteome Discoverer with the Precursor Ions Quantifier workflow using Minora feature detection.

For protein-detection counts, proteins with Number of Peptides ≥ 2 were retained. For principal component analysis (PCA), proteins with Number of Peptides ≥ 2 that were quantified in all 12 samples were used. Abundance values were log2-transformed, and each sample was median-normalized by shifting its median to the median across the 12 sample-specific medians. PCA was performed in R using the prcomp function with centering and without scaling.

Differential protein abundance was analyzed in Perseus v1.6.15.0 [22]. Protein abundance values were transformed to log2 scale, and each sample was median-centered to reduce sample-level intensity offsets. Missing entries were replaced using the standard Perseus imputation settings applied here: Gaussian width = 0.3 and downshift = 1.8. Pairwise differences in protein abundance were evaluated using Student’s t-tests in Perseus. Proteins were considered differentially changed based on a nominal p value < 0.05, |log2 fold change| ≥ log2(1.5), and Number of Peptides ≥ 2. No multiple-testing correction was applied to the primary differential-change criteria. The three comparisons used for this analysis were single CB versus control, repeated CB versus control, and repeated CB versus single CB. Differentially changed proteins are reported in Supplementary Table 1.

To compare single and repeated CB responses, the union of differentially changed proteins from the three pairwise comparisons was classified into response classes based on differential-change patterns, directionality, and relative response magnitude. For classification based on relative response magnitude, the absolute response was defined as the absolute log2 fold change relative to the control group. Repeated-specific proteins were defined as proteins that met the differential-change criteria in repeated CB-treated muscles compared with control muscles but did not meet these criteria in single CB-treated muscles compared with control muscles. Single-versus-repeated difference-only proteins were defined as proteins that met the differential-change criteria in the repeated CB versus single CB comparison but did not meet these criteria in either comparison with the control group. Sustained proteins were defined as proteins that met the differential-change criteria in both comparisons with the control group and changed in the same direction, with the absolute response to repeated CB ranging from 0.75-fold to 1.25-fold of the absolute response to single CB. Reinforced proteins were defined as proteins that met the differential-change criteria in both comparisons with the control group and changed in the same direction, with the absolute response to repeated CB greater than 1.25-fold of the absolute response to single CB. Single-specific/weakly attenuated proteins were defined as proteins that met the differential-change criteria in single CB-treated muscles compared with control muscles but did not meet these criteria in repeated CB-treated muscles compared with control muscles, with the absolute response to repeated CB remaining at least 0.75-fold of the absolute response to single CB. Attenuated to baseline proteins were defined as proteins that met the differential-change criteria in single CB-treated muscles compared with control muscles but did not meet these criteria in repeated CB-treated muscles compared with control muscles, with the absolute response to repeated CB reduced to less than 0.75-fold of the absolute response to single CB. Attenuated proteins were defined as proteins that met the differential-change criteria in both comparisons with the control group and changed in the same direction, with the absolute response to repeated CB less than 0.75-fold of the absolute response to single CB. Reversed proteins were defined as proteins that met the differential-change criteria in both comparisons with the control group but changed in opposite directions. Proteins that did not match these defined patterns were assigned to an unclear-response category. Response-class assignments are provided in Supplementary Table 2.

Protein-count, PCA, and volcano plots were generated in R using the ggplot2 package, with ggrepel used for sample or protein labels. Response-class heatmaps were generated using the pheatmap package, and the two heatmap panels were combined using the gridExtra package. For heatmap visualization, sample-level protein abundance values were converted to row-wise z-scores. Proteins and samples were not hierarchically clustered; proteins were grouped according to their response-class assignments.

### Immunoblotting

Selected proteins highlighted by the proteomic analysis were examined by immunoblotting. Plantaris muscles were extracted in the same RIPA buffer containing protease inhibitors as described above. Homogenates were centrifuged at 15,000 × g for 30 min at 4°C, and the supernatants were used as cleared lysates. Protein amounts were measured with the BCA Protein Assay Kit (Takara).

Equivalent amounts of total protein were resolved by SDS-PAGE and electrotransferred to PVDF membranes. Membranes were blocked in skim milk/TBST and then reacted overnight at 4°C with primary antibodies. The antibodies used were anti-Klhl40 (1/2000, 21836-1-AP, Proteintech), anti-Napa (1/2000, 10546-1-AP, Proteintech), anti-Galectin-3 (1/2000, AF1197, R&D Systems, Inc.), anti-Rab18 (1/2000, 11304-1-AP, Proteintech), and anti-α-tubulin (1/2000, #2144, CST, used with Can Get Signal reagent; Toyobo). HRP-conjugated anti-rabbit IgG (1/10000, #7074, CST) and anti-goat IgG (H+L) (1/10000, 81-1620, Thermo Fisher Scientific) served as secondary antibodies. Chemiluminescent detection was performed with ImmunoStar LD reagent (FUJIFILM Wako Pure Chemical), and images were acquired with a cooled CCD imaging system (Light-Capture, ATTO). Band density was measured in Fiji ver. 2.16.0 and normalized to α-tubulin. Full-length immunoblot images are shown in Supplementary Figure 2.

## Results

### Repeated intermittent CB exposure produced a distinct muscle-weight response under a long-term treatment schedule

We first examined whether the increase in muscle mass induced by a 2-week CB treatment persisted after treatment cessation. Mice were treated with or without CB for 2 weeks, and muscles were collected either immediately after this treatment period or after an additional 4-week period without CB. Control and CB-treated groups were included at both collection time points. Body weight did not differ significantly between control and CB-treated mice at either time point (Supplementary Fig. 1A). At the end of the 2-week treatment period, CB significantly increased Gast and plantaris weights compared with the corresponding control groups, whereas these differences were no longer significant after the additional 4-week period without CB (Supplementary Fig. 1B,C). Soleus weight showed an increasing trend after the 2-week CB treatment, but this change did not reach statistical significance (Supplementary Fig. 1D). These results indicated that the muscle-weight effect induced by a 2-week CB exposure was substantially reduced after a 4-week post-treatment period.

Based on this observation, we designed a repeated intermittent CB protocol to compare control, single CB, and repeated CB groups under the same long-term experimental timeline (Fig. 1A). Muscles were collected at the end of the final 2-week treatment period. Body weight did not significantly differ among the three groups (Fig. 1B). In contrast, repeated CB mice showed significantly increased Gast, plantaris, and soleus weights compared with control mice (Fig. 1C–E). The single CB-treated group showed a trend toward modest increases in some muscle weights. In addition, soleus weight was significantly higher in repeated CB mice than in single CB mice (Fig. 1E). These results suggest that the muscle-weight response to the final 2-week CB exposure alone was relatively limited under this long-term treatment schedule, whereas repeated intermittent CB exposure produced a more evident muscle-weight response.

### Proteomic profiling revealed different global patterns after single and repeated CB exposure

To determine whether the different muscle-weight responses were accompanied by changes in protein abundance, we performed proteomic analysis of plantaris muscles from control, single CB, and repeated CB mice. Protein-detection depth was comparable among samples after peptide-based filtering (Fig. 2A). PCA of commonly quantified proteins showed that single CB samples had a more dispersed distribution, whereas repeated CB samples formed a more compact cluster (Fig. 2B). Thus, the plantaris proteome after repeated CB exposure showed a global pattern that was not identical to that observed after a single CB exposure.

**Figure 2.**
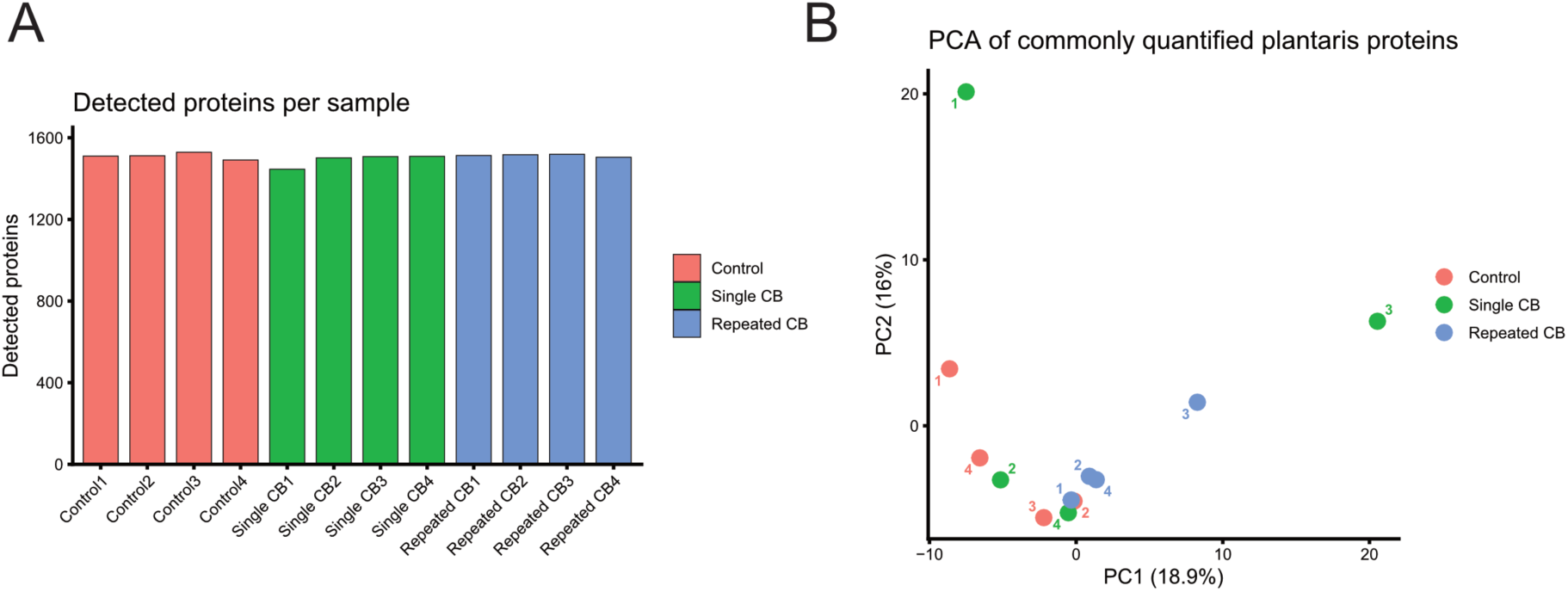
Proteomic overview of plantaris muscle after single and repeated CB exposure. (A) Number of detected proteins per sample in plantaris muscles from control, single CB, and repeated CB mice after filtering for Number of Peptides ≥ 2. (B) PCA of commonly quantified plantaris proteins after the same filtering. Each point represents one biological sample, and numbers indicate individual sample indices within each group. Data are from n = 4 mice per group.

### Single and repeated CB exposure induced overlapping but distinct proteomic changes

We next compared protein abundance between groups using the criteria described in Methods. In the single CB versus control comparison, 49 proteins were significantly changed, including 23 increased and 26 decreased proteins (Fig. 3A and Supplementary Table 1). In the repeated CB versus control comparison, 74 proteins were significantly changed, including 47 increased and 27 decreased proteins (Fig. 3B and Supplementary Table 1). Thus, repeated CB exposure altered a larger number of proteins relative to control than did a single CB exposure.

**Figure 3.**
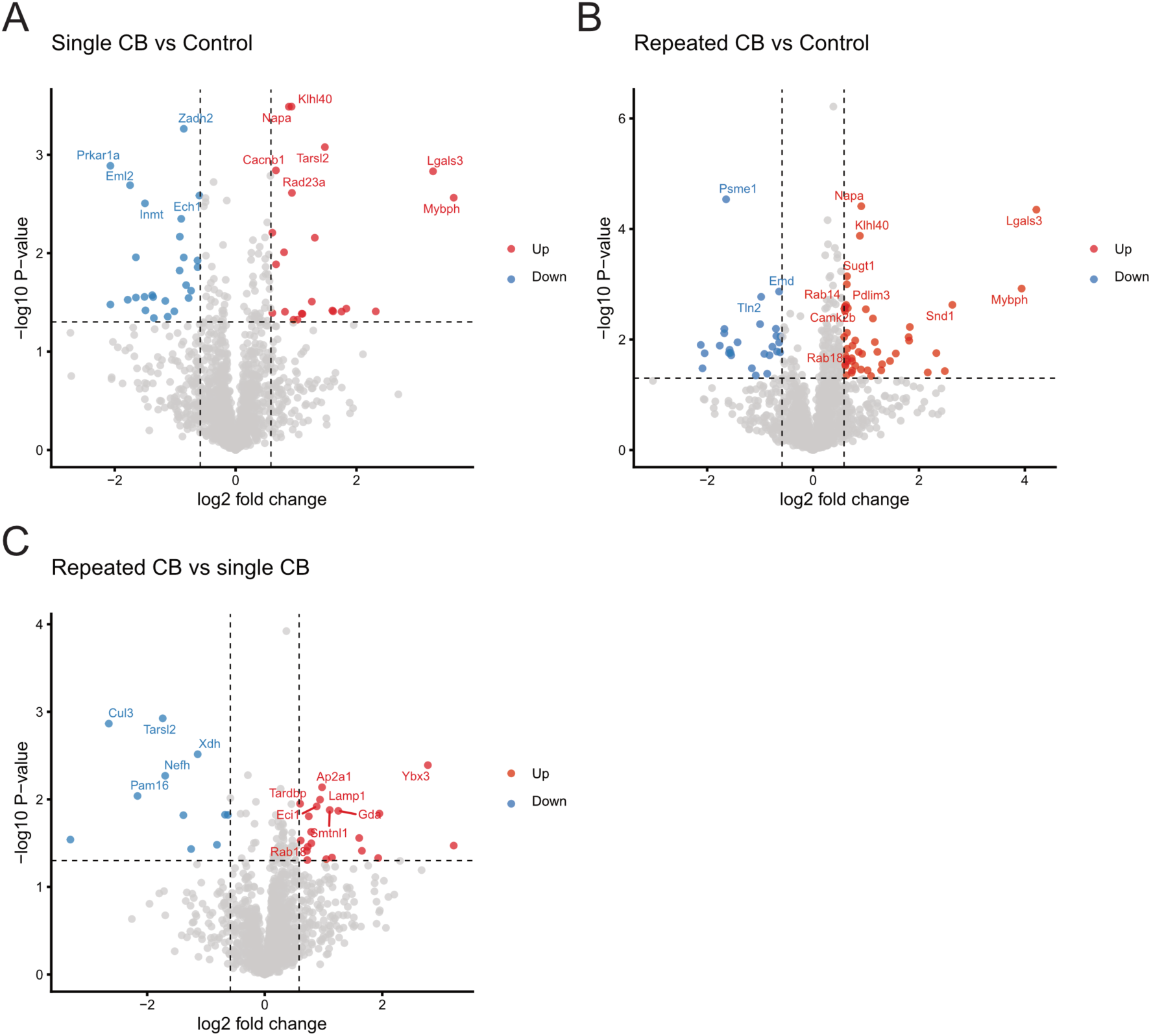
Differential proteomic responses to single and repeated CB exposure. Volcano plots showing differentially changed proteins in plantaris muscle. Comparisons are shown for single CB versus control (A), repeated CB versus control (B), and repeated CB versus single CB (C). Proteins passing the peptide filter described in Methods were analyzed. Red and blue points indicate significantly increased and decreased proteins, respectively, based on p < 0.05 and |log2 fold change| ≥ log2(1.5). Differentially changed proteins from each comparison are listed in Supplementary Table 1.

A direct comparison between repeated CB and single CB muscles identified 32 significantly changed proteins, including 21 increased and 11 decreased proteins in the repeated CB group (Fig. 3C and Supplementary Table 1). These results indicate that repeated CB exposure does not simply reproduce the proteomic response induced by a single CB exposure.

### Response-class analysis identified multiple patterns of CB-associated proteomic remodeling

To further define the relationship between single CB and repeated CB responses, we integrated the three pairwise comparisons and classified proteins into response classes (Supplementary Table 2). This classification was based on whether each protein changed after single CB exposure, after repeated CB exposure, and between repeated CB and single CB muscles.

Proteins were assigned to response classes according to the criteria described in Methods. Among the 130 proteins included in the heatmap, 57 were classified as repeated-specific, 24 as single-versus-repeated difference-only, 13 as sustained, 2 as reinforced, 6 as single-specific/weakly attenuated, 26 as attenuated to baseline, 1 as attenuated, and 1 as reversed (Fig. 4 and Supplementary Table 2). Thus, proteins whose changes became apparent after repeated exposure represented the largest defined response class.

**Figure 4.**
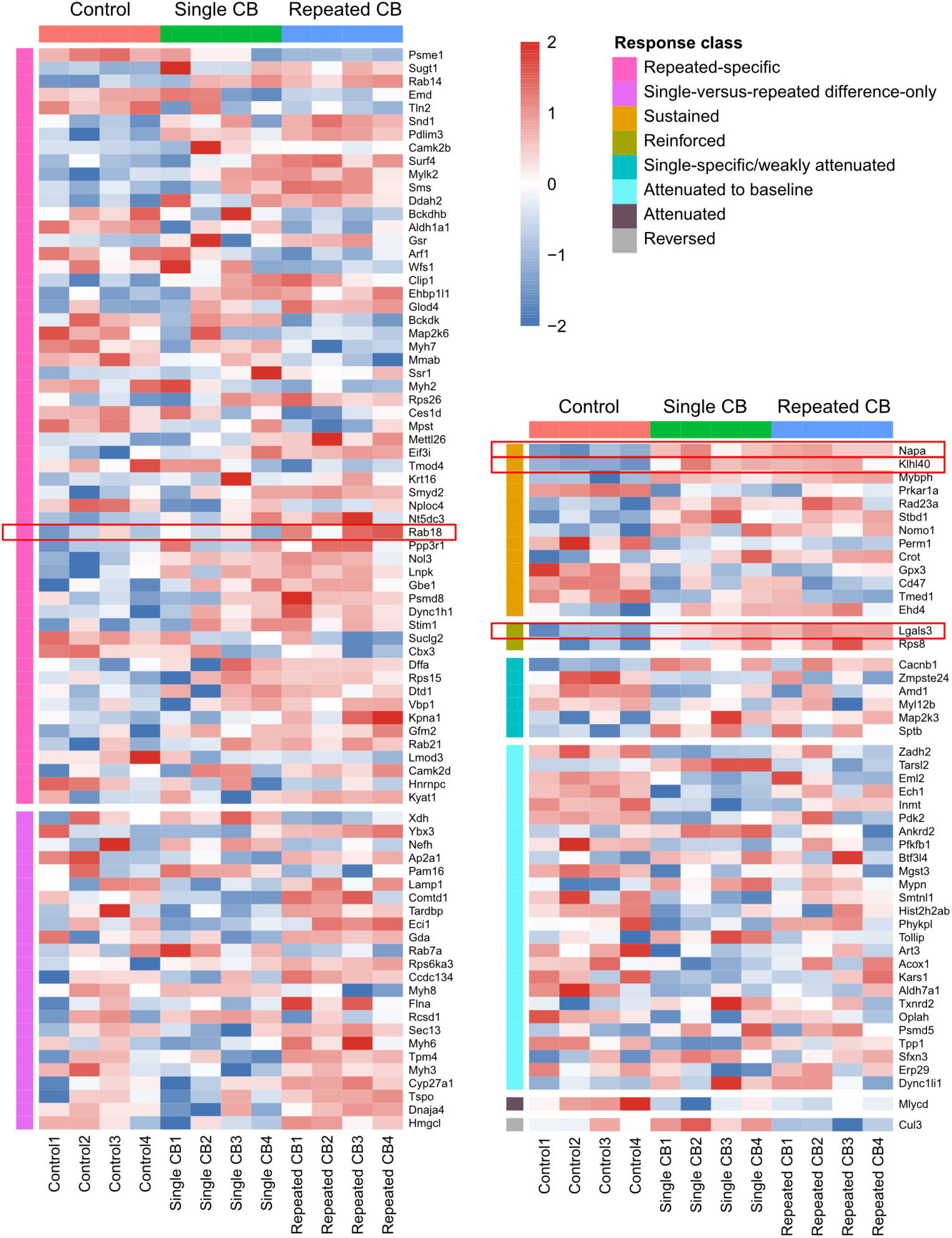
Response-class heatmap of CB-associated proteomic changes. The heatmap shows all 130 response-classified proteins identified by integrated analysis of the three pairwise comparisons, excluding proteins assigned to the unclear-response category. Protein abundance values are shown as row-wise z-scores across control, single CB, and repeated CB samples, with values truncated at −2 and +2 for visualization. Samples are arranged by group, and proteins are grouped according to their response-class assignment. The two heatmap panels group related response classes: repeated-specific and single-versus-repeated difference-only proteins are shown together, whereas sustained, reinforced, single-specific/weakly attenuated, attenuated to baseline, attenuated, and reversed proteins are shown together. Response-class assignments are provided in Supplementary Table 2. Proteins selected for immunoblot validation are highlighted by red boxes.

The heatmap revealed multiple abundance patterns across control, single CB, and repeated CB muscles. The repeated-specific class formed the largest group, indicating that many proteins met the predefined differential-change criteria only in the repeated CB-versus-control comparison. The single-versus-repeated difference-only class comprised proteins that did not meet the differential-change criteria in either comparison with the control group but met these criteria in the direct comparison between single CB and repeated CB muscles. The sustained and reinforced classes represented protein responses shared by single and repeated CB exposure, whereas the single-specific/weakly attenuated, attenuated to baseline, and attenuated classes represented responses that were more prominent after single CB exposure or were reduced after repeated exposure. The reversed class contained proteins that met the differential-change criteria after both single and repeated CB exposure but changed in opposite directions. These results indicate that the CB-induced proteomic response consists of multiple response patterns rather than a uniform progression from the single-exposure response to the repeated-exposure response.

### Immunoblotting validated selected response-class proteins

Finally, we performed immunoblotting to validate selected proteins identified by the response-class analysis. Proteins were selected to represent distinct biological categories within the response-classified proteome, while also considering response pattern, clarity of the proteomic change, and antibody availability. We selected Klhl40 as a sustained sarcomere-associated protein, Napa as a sustained membrane-trafficking-related protein, Lgals3/Galectin-3 as a tissue-remodeling-related protein classified as reinforced in the proteomic analysis, and Rab18 as a protein classified as repeated-specific in the proteomic analysis (Fig. 5A).

**Figure 5.**
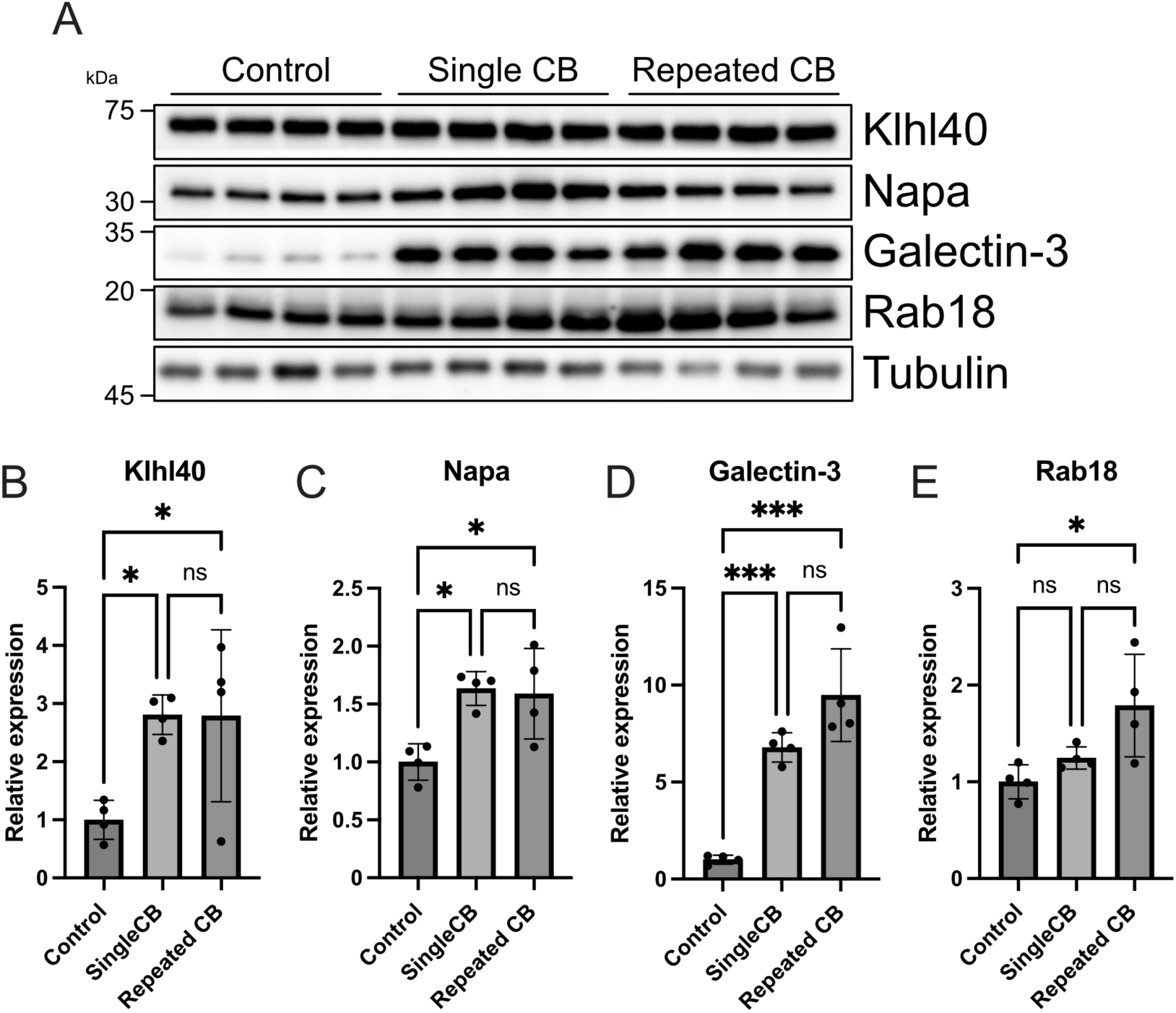
Immunoblot validation of selected response-class proteins. (A) Representative immunoblots for Klhl40, Napa, Lgals3/Galectin-3, Rab18, and α-tubulin in plantaris muscles from control, single CB, and repeated CB mice. (B–E) Quantification of Klhl40 (B), Napa (C), Lgals3/Galectin-3 (D), and Rab18 (E) protein abundance. Band intensities were normalized to α-tubulin and expressed relative to the control group. Data are presented as mean ± s.d.; n = 5 mice per group. Data were analyzed by ordinary one-way ANOVA followed by Tukey’s multiple-comparison test. ns, not significant; *P < 0.05; ***P < 0.001. Uncropped immunoblots are provided in Supplementary Figure 2.

Immunoblotting showed that Klhl40 was increased in both single CB and repeated CB muscles compared with control muscles, with no significant difference between the two CB groups (Fig. 5B). Napa showed a similar pattern, increasing in both CB-treated groups relative to control (Fig. 5C). Galectin-3 showed a particularly marked increase after CB exposure and was increased in both single CB and repeated CB muscles (Fig. 5D). Rab18 did not significantly differ between control and single CB muscles, but was increased in repeated CB muscles compared with control muscles (Fig. 5E). These immunoblotting results confirmed representative proteomic changes identified by the response-class analysis and supported the presence of shared, strongly CB-responsive, and repeated CB-associated protein responses in plantaris muscle.

## Discussion

In this study, we established a repeated intermittent CB administration model consisting of 2-week CB treatment periods separated by 4-week washout periods, and examined the effects of single and repeated CB exposure on mouse skeletal muscle using label-free quantitative proteomics. PCA showed that the single CB group exhibited a relatively dispersed distribution, whereas the repeated CB group formed a more compact cluster. In addition, the sets of significantly altered proteins differed partially among the single CB vs control, repeated CB vs control, and repeated CB vs single CB comparisons. Furthermore, response-class analysis integrating these three pairwise comparisons revealed that the repeated-specific class represented the largest group among clearly classified proteins, indicating that many proteins met the predefined differential-change criteria only in the repeated CB-versus-control comparison. These results suggest that the skeletal muscle response to repeated CB administration is not merely a reiteration of the single-exposure response, but is accompanied by exposure history-dependent changes in the proteomic state.

Recent MS-based proteomic analysis of human skeletal muscle has shown that prolonged β2-adrenergic stimulation with terbutaline increases kelch-like family proteins, including KLHL40 and KLHL41 [17]. In addition, oral CB, but not therapeutic inhaled formoterol, has been reported to increase KLHL41 abundance in human skeletal muscle [23]. In the present study, Klhl40 protein was increased in both the single CB and repeated CB groups. Klhl40 is a skeletal muscle-enriched kelch-like protein, and previous studies have shown that Klhl40 deficiency destabilizes thin filament proteins and causes nemaline-like myopathy, indicating its importance in maintaining sarcomere structure in skeletal muscle [24]. The increase in Klhl40 observed here suggests that CB exposure alters responses related to the contractile apparatus or myofibrillar protein homeostasis in plantaris muscle. Consistent with this interpretation, our proteomic analysis also showed a marked increase in Mybph in both the single CB and repeated CB groups, suggesting that sarcomere-associated proteins are altered after CB exposure. Together, these findings support the possibility that kelch-like family proteins constitute part of a sarcomere-associated proteomic response to prolonged or repeated β2-adrenergic stimulation.

Galectin-3 is a β-galactoside-binding lectin involved in diverse tissue-remodeling processes, including inflammation, immune responses, cell migration, tissue repair, and fibrosis [25,26]. In the present study, Galectin-3 was markedly increased in both the single CB and repeated CB groups, with a stronger trend in the repeated CB group, although the difference between single CB and repeated CB was not statistically significant. This result suggests that CB-treated muscle undergoes not only changes in myofiber-associated proteins but also alterations in the tissue environment, including inflammatory, immune, or interstitial remodeling responses. Tissue-wide remodeling involving immune cells, stromal cells, blood vessels, and extracellular matrix components has been reported during muscle hypertrophy [27]. Interestingly, in our proteomic dataset, broad increases in typical extracellular matrix remodeling-related proteins, such as collagen family proteins and Serpinh1, were not clearly observed. Moreover, the present analysis cannot determine whether the increased Galectin-3 was derived from myofibers or from non-myogenic cells such as macrophages or interstitial cells. Therefore, further localization analyses will be required to clarify whether Galectin-3 primarily reflects inflammatory or immune-cell responses, interstitial remodeling, or a myofiber-associated response.

Napa is an alpha-soluble NSF attachment protein (α-SNAP) that functions together with NSF to regulate SNARE complex disassembly and is involved in membrane fusion and vesicular trafficking [28,29]. In this study, Napa was increased in both the single CB and repeated CB groups, indicating that SNARE/NSF-related membrane trafficking proteins are altered after CB exposure in skeletal muscle. Rab18 is a small GTPase implicated in lipid droplets, ER membrane dynamics, and ER–lipid droplet contact sites [30,31]. In our proteomic analysis, Rab18 was classified as a repeated-specific protein and was significantly increased in the repeated CB group compared with the control group by immunoblotting. However, because immunoblotting did not detect a statistically significant difference between the single CB and repeated CB groups, Rab18 should be interpreted as a candidate endomembrane/lipid droplet-related protein change that becomes more clearly detectable under repeated CB conditions, rather than as a strictly repeated CB-specific factor. In addition to Rab18, the proteomic analysis identified changes in other membrane trafficking or endomembrane-related proteins, including Rab14, Rab21, Lnpk, and Lamp1. These changes may represent a characteristic feature of the plantaris muscle proteome response to repeated CB exposure. Taken together, the changes in Napa and Rab18 suggest that, in addition to sarcomere-associated and tissue-remodeling-related proteins, proteins involved in membrane fusion, vesicular trafficking, and lipid droplet/ER-associated membrane dynamics are also altered after CB exposure.

A previous human fiber type-specific proteomic study also reported that β2-adrenergic stimulation increases ribosomal proteins and decreases mitochondrial electron transport chain proteins in both type I and type II muscle fibers [32]. In the present study, the repeated CB group also showed increases in a subset of ribosomal or translation-related proteins and decreases in several metabolic or mitochondrial proteins. Therefore, although these pathways were not the primary focus of the present analysis, these findings in mouse plantaris muscle partially overlap with proteomic changes reported in human skeletal muscle after β2-adrenergic stimulation.

Overall, this study demonstrates that single and repeated intermittent CB administration induce both shared and repeated exposure-associated proteomic responses in skeletal muscle. Although repeated CB administration produced a more evident muscle-weight response, the underlying molecular changes were not a simple amplification of the single CB response. Instead, repeated CB exposure was associated with complex proteome remodeling involving sarcomere- and myofibril-related responses, translation- and metabolism-related responses, tissue-remodeling-related responses, and membrane-trafficking-related responses. CB is a prohibited doping substance. β2-Adrenergic agonists induce bronchodilation through relaxation of airway smooth muscle and can also affect skeletal muscle mass and metabolism. Therefore, understanding how repeated CB exposure alters the skeletal muscle proteome is important not only for clarifying the molecular basis of β2-adrenergic stimulation-induced muscle hypertrophy, but also for evaluating the biological consequences of repeated CB exposure. Thus, the present dataset provides a foundation for future mechanistic studies of CB-induced skeletal muscle hypertrophy and remodeling associated with repeated exposure.

## Institutional Review Board Statement

Animal procedures were authorized by the Fujita Health University Institutional Animal Care and Use Committee, Japan (approval No. APU25120-MD1), and were performed under the relevant institutional animal-care rules.

## Informed Consent Statement

Not applicable.

## Data Availability and Statistical Reporting

Values are shown as mean ± s.d. GraphPad Prism 9 was used for statistical analyses of body weight, muscle weight, and immunoblot quantification. In Fig. 1, body weight and muscle-weight data from the repeated intermittent CB experiment were evaluated by ordinary one-way ANOVA followed by Tukey’s multiple-comparison test. In Supplementary Fig. 1, data from the 2-week treatment and 4-week post-treatment experiment were analyzed by ordinary two-way ANOVA using treatment and time point as factors, followed by Šídák’s multiple-comparison test for control-versus-CB comparisons within each time point. Immunoblot quantification across the control, single CB, and repeated CB groups was analyzed by ordinary one-way ANOVA followed by Tukey’s multiple-comparison test. For these analyses, statistical significance was set at p < 0.05.

For the proteomic analyses, pairwise differences in protein abundance were evaluated using Student’s t-tests in Perseus. Proteins were considered differentially changed based on a nominal p value < 0.05, |log2 fold change| ≥ log2(1.5), and Number of Peptides ≥ 2. No multiple-testing correction was applied to the primary differential-change criteria. The figure legends specify the sample sizes and statistical tests used for each dataset.

Supplementary Table 1 contains the proteins identified as differentially changed in the three pairwise proteomic comparisons. Supplementary Table 2 provides the response-class assignments for all 130 proteins included in the response-class heatmap. Supplementary Figure 2 includes the uncropped immunoblot images. The LC-MS/MS raw files generated in this study have been deposited in the ProteomeXchange Consortium [33] via jPOST [34] under the dataset identifiers PXD080839 and JPST004754. Other data supporting the conclusions of this study are available from the corresponding author upon reasonable request.

## Author Contributions

Conceptualization, K.H.; methodology, K.H. and H.Y.; investigation, K.H., H.Y., Y.K., and K.T.; formal analysis, K.H. and H.Y.; data curation, K.H. and H.Y.; writing—original draft preparation, K.H.; writing—review and editing, K.H., H.Y., Y.K., and K.T.; supervision, K.H. and K.T.; funding acquisition, K.H. and K.T. All authors have read and agreed to the submitted version of the manuscript.

## Acknowledgements

We gratefully acknowledge E. Fukushima, S. Takeuchi, C. Ohshima, and the staff of Fujita Health University Open Facility Center for experimental assistance.

## Declaration of AI-assisted technologies in manuscript preparation

ChatGPT (OpenAI) was used only for English editing and organization of the manuscript text. All scientific content, data presentation, and interpretation were checked and finalized by the authors.

## Supplementary Materials

### Supplementary figures and figure legends

**Supplementary Figure 1.**
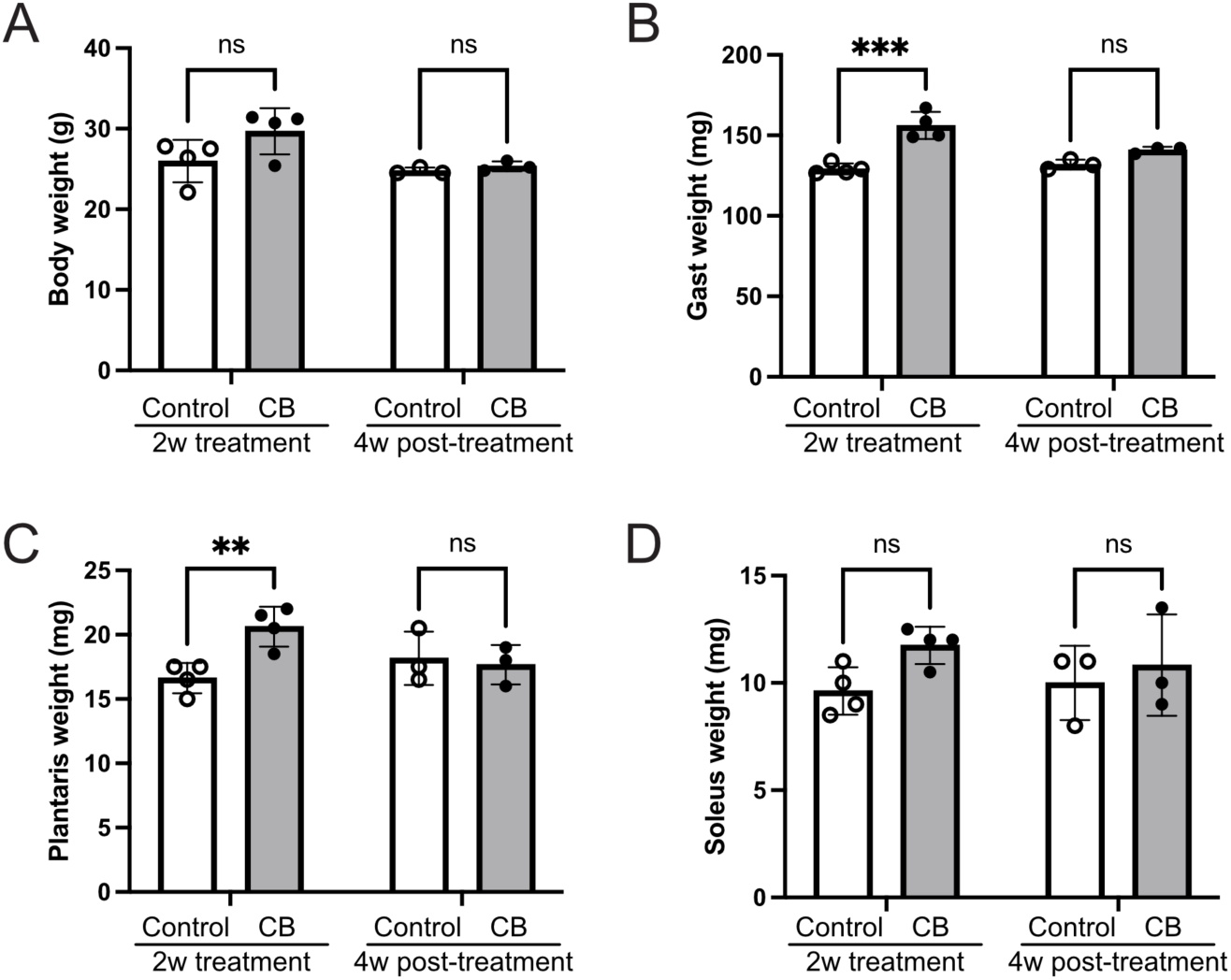
Muscle-weight changes after 2-week CB treatment and a 4-week post-treatment period. Body weight (A), Gast weight (B), plantaris weight (C), and soleus weight (D) were measured after mice were treated with or without CB in the drinking water for 2 weeks. Measurements were performed either immediately after the 2-week treatment period or after an additional 4- week period without CB. Data are presented as mean ± s.d.; n = 4 mice per group at the 2-week treatment time point and n = 3 mice per group at the 4-week post-treatment time point. Data were analyzed by ordinary two-way ANOVA with treatment and time point as factors, followed by Šídák’s multiple-comparison test comparing control and CB-treated groups within each time point. ns, not significant; **P < 0.01; ***P < 0.001.

**Supplementary Figure 2.**
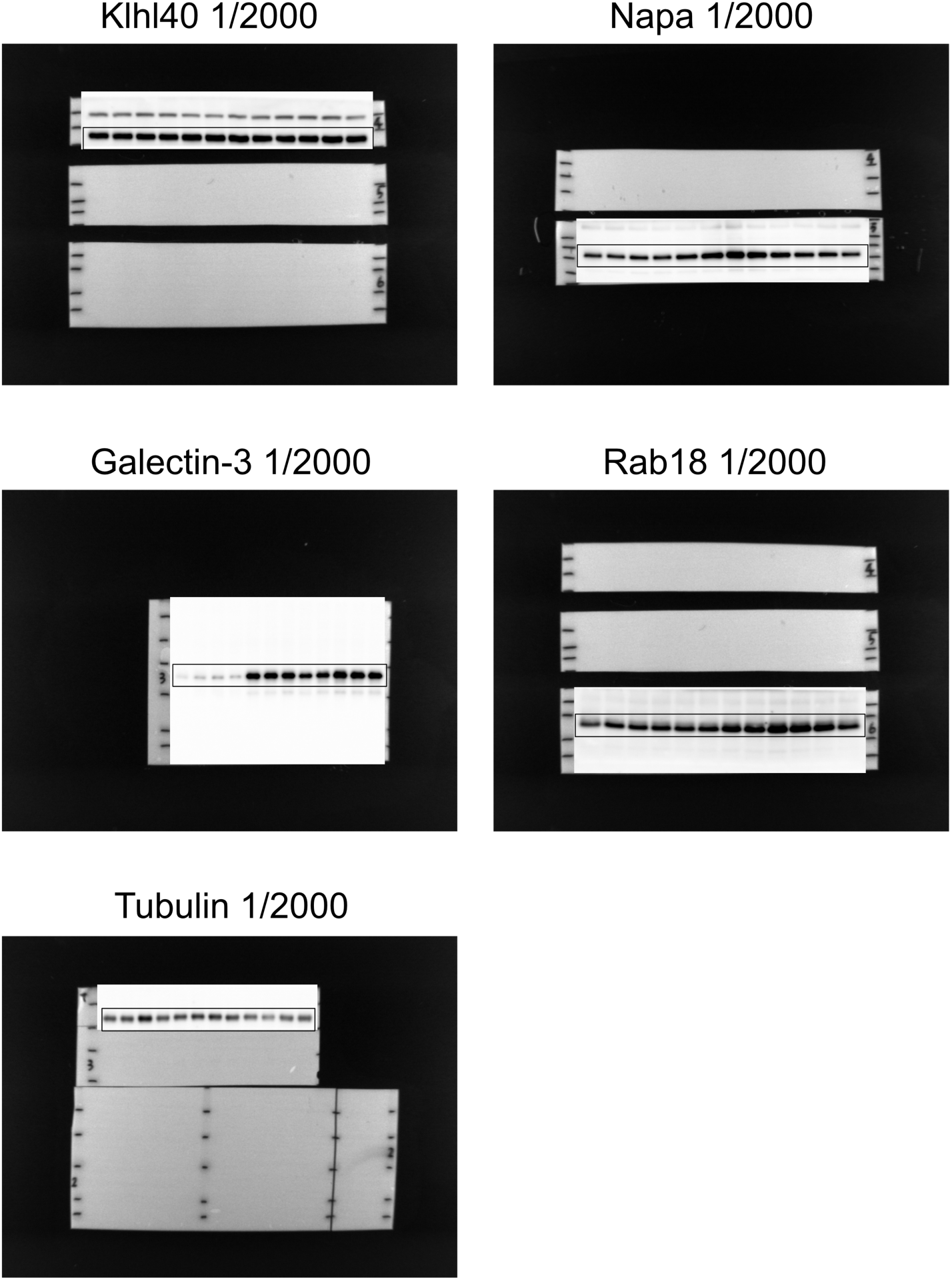
Uncropped immunoblots for Fig. 5. Uncropped immunoblot images corresponding to the cropped blots shown in Fig. 5. Antibody names and dilutions are indicated in each panel.

**Supplementary Table 1.** Differentially changed proteins identified in the three pairwise proteomic comparisons.

**Supplementary Table 2.** Response-class assignments for all 130 proteins displayed in the response-class heatmap.

